# Conserved Roles of Sp1 in Zebrafish Development and Early Organogenesis

**DOI:** 10.64898/2026.01.12.699150

**Authors:** Ankita Sharma, Sudiksha Mishra, Greg Jude Dsilva, Saurabh J Pradhan, Pavan Dev Govardhan, Sanjeev Galande

## Abstract

Transcription factors (TFs) function in combinatorial networks to orchestrate gene expression during development and disease. Yet, the mechanisms underlying their context-specific regulatory roles remain poorly understood. Here, we investigate Specificity Protein 1 (Sp1), the first sequence-specific transcription factor identified and a key regulator of TATA-less gene promoters as a model to elucidate conserved and lineage-specific functions in vertebrate development. While Sp1 is extensively studied in mammalian systems, particularly in cancer progression, its evolutionary acquisition and developmental roles in early vertebrates remain unexplored. Using zebrafish as a model, we characterized the spatiotemporal expression dynamics and functional significance of *sp1* during embryogenesis. Loss-of-function analyses revealed profound morphological defects across multiple organ systems, including skeletal, neural, and gastrointestinal tissues. Complementary stage-specific transcriptomic profiling uncovered widespread dysregulation of lineage-defining genes and perturbations in core cellular pathways related to the cell cycle, DNA metabolism, and apoptosis. Together, our findings establish Sp1 as a pivotal transcriptional regulator in early vertebrate development and reveal mechanistic insights into how conserved TF networks coordinate developmental programs and context-dependent disease states.

**Summary statement:** This study characterizes the expression profile and establishes the conserved role of SP1 in specific processes during zebrafish development, which were previously unreported.

## Introduction

Developmental processes rely on tightly coordinated gene expression, governed by networks of transcription factors (TFs) that define cell identity, drive cellular differentiation, tissue morphogenesis, and organ development. By integrating diverse regulatory inputs and interaction with co-factors, TFs ensure temporal and spatial control of transcriptional programs essential for normal physiology. Dysregulation of these factors, however, is frequently associated with developmental abnormalities and a wide spectrum of diseases, including cancer and metabolic disorders. Despite their central importance, the mechanisms by which TFs achieve context-specific regulation responding to developmental cues, cellular states, or environmental signals remain incompletely understood. Deciphering how functions of such TFs evolved during evolution with a conserved function in lineage-specific transcriptional networks is therefore fundamental to understanding the logic of gene regulation during vertebrate development and disease.

Specificity protein 1 (SP1) was the first transcription factor to be identified, originally isolated from HeLa cells for its ability to bind GC boxes and activate transcription from the SV40 early and thymidine kinase promoters^1–3^. By recognizing GC-rich regions of the genome, SP1 drives expression of many genes that lack conventional TATA box-containing promoters^2^. It belongs to the SP family of C2H2 zinc finger transcription factors, within the larger Sp/Krüppel-like factor (KLF) family^4^. SP1 provides a compelling model for uncovering how broadly expressed transcription factors achieve regulatory specificity. Although ubiquitously expressed and traditionally viewed as a “general” regulator of housekeeping and cell-growth genes, SP1 exhibits highly context-dependent activity shaped by chromatin state, combinatorial interactions with other TFs, and extensive post-translational modifications. Functionally, SP1 is required for diverse processes, including proliferation, differentiation, apoptosis, angiogenesis, and migration, and is essential for normal development and adult homeostasis^5–15^. Dysregulation of SP1 has been implicated in various diseases, notably cancer and neurological disorders such as Huntington’s and Alzheimer’s disease^16–19^. Studying SP1 during embryogenesis therefore offers an opportunity to dissect how a widely expressed TF integrates environmental and developmental cues to orchestrate lineage-specific transcriptional programs, serving as an exemplar for understanding gene regulatory logic more broadly.

Despite extensive work in human cell lines and mouse models, the function of SP1 during vertebrate development has been less well explored. A few zebrafish orthologs of certain Sp family members, such as *sp2* and *sp5-like*, have been cloned and shown to play roles in early development^20–22^. Zebrafish Sp1 itself has been purified and shown to possess DNA-binding activity^23^ and limited phenotypic observations related to retinal development have been described^24^. However, its expression dynamics and developmental functions in zebrafish have not been comprehensively explored. Here, we investigated the role of Sp1 in zebrafish embryogenesis. We demonstrate that *sp1* is expressed ubiquitously in early development, before becoming enriched in the anterior region and gastrointestinal system. Loss-of-function analysis resulted in developmental abnormalities during larval stages, underscoring its essential role. Transcriptome profiling revealed dysregulation of genes associated with the cell cycle, DNA metabolism, regeneration, neuronal development, and eye formation. Together, our findings establish that zebrafish Sp1 performs conserved fundamental functions while providing a tractable model to study its developmental roles in vivo.

## Results

### The functional domains of Sp1 show sequence conservation across species

We investigated the evolutionary conservation of SP1 across the vertebrates. Phylogenetic analysis of SP1 orthologs across diverse taxa, including *Danio rerio, Homo sapiens, Mus musculus, Rana temporaria, Rattus norvegicus, Bos taurus, Macaca mulatta, Gallus gallus, Pan troglodytes, Oryzias latipes, Alligator mississippiensis, Sus scrofa, Taeniopygia guttata, Zootoca vivipara, Trachemys scripta elegans, Latimeria chalumnae, Protopterus annectens* and *Branchiostoma belcheri*, highlighted the evolutionary divergence between zebrafish and human SP1 proteins (Figure 1A). This divergence likely reflects both faster sequence evolution in the teleost lineage and the teleost-specific whole genome duplication (TSGD)^25^. Based on the latest genome assembly (GCA_000002035.4), the updated zebrafish *sp1* transcript (NM_212662.2) and protein (NP_997827.2) are 2312 bp and 594 aa, respectively. Pairwise analysis showed ∼50% identity between human and zebrafish SP1 (Suppl Figure S1A), consistent with previous reports (∼49.5%)^23^.

**Figure 1.**
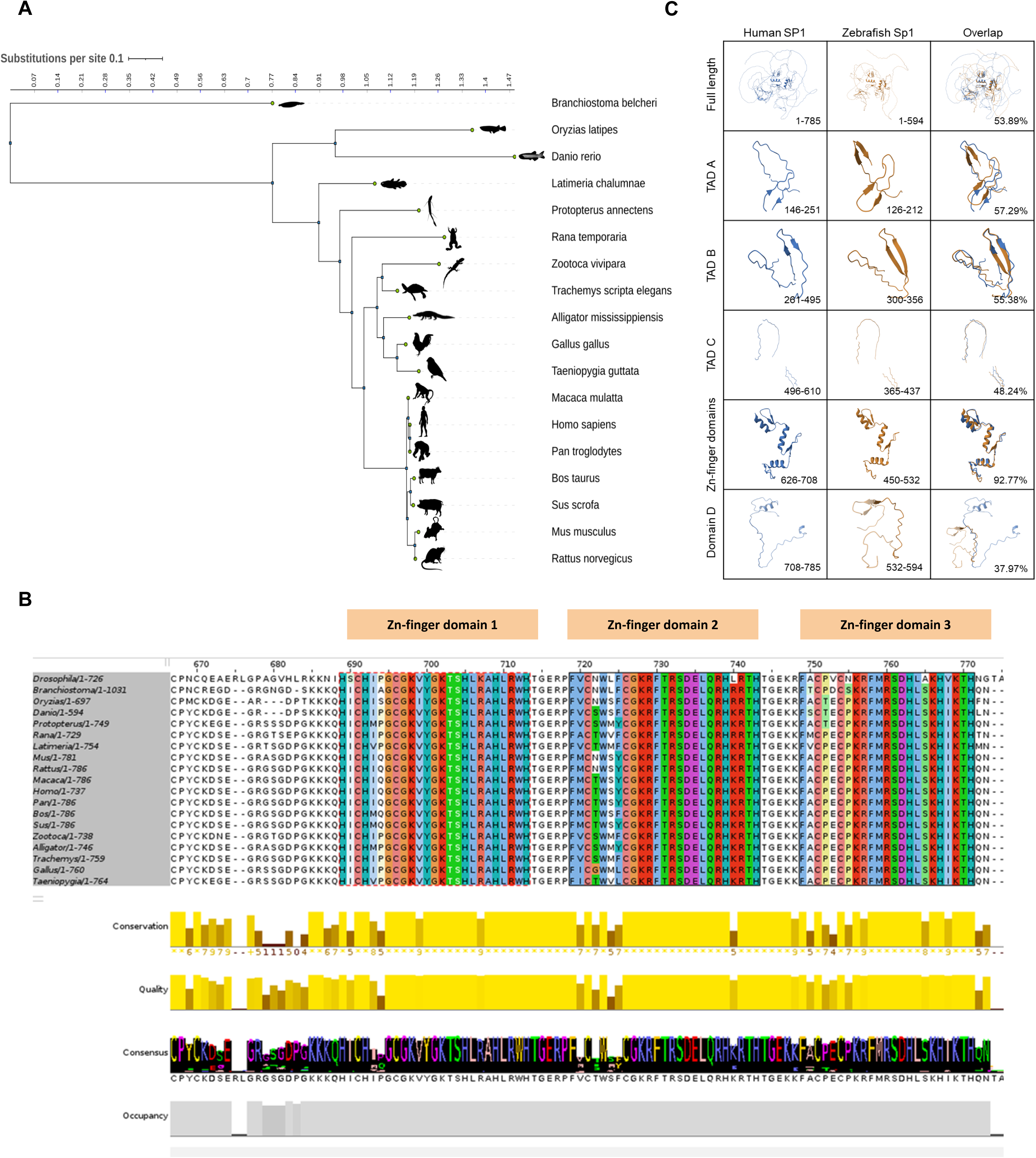
Phylogenetic analysis of SP1 shows conservation across evolution. **A.** Phylogenetic tree of SP1 orthologs among the species indicated in the schematic using the RAxML-NG algorithm on CIPRES. The tree scale of 1 corresponds to 1 substitution per site. **B.** Multiple sequence alignment of the Zn-finger domains for SP1 orthologs using Clustal Omega. **C.** Predicted TADs, Zn-finger domains and Domain D of zebrafish Sp1 aligned with the corresponding domains of the human SP1 proteins. The lower right side of each box (left to right) shows the amino acid residues reported for each Human SP1 domain, predicted residues for Zebrafish Sp1 domains, and percent similarity between the two corresponding amino acid stretches in the overlap.

Further, we asked whether conservation was enriched within the functional domains. Multiple sequence alignment^26^ as well as domain prediction and alignment of SP1 orthologs through HMMER^26^ using the Pfam database confirmed that the zinc finger domains are conserved and localized at the C-terminal across species (Figure 1B & Figure S1B). Additionally, the zinc finger domains of zebrafish Sp1 were rich in glutamine residues, a characteristic shared with the zinc finger domains of human SP1^27^ (Figure 1C). We then aligned the full zebrafish Sp1 peptide with all the reported human SP1 domains to identify putative homologous regions other than the zinc finger regions. Human SP1 has three C-terminal zinc finger domains essential for DNA binding, a repressor domain, three transactivation domains (TAD-A, –B, and –C), and Domain D, which is required for oligomerisation and synergistic transcriptional activation^28–30^. We were able to identify the homologous regions for most domains, though with variable identity (37–55%) (Figure 1C) and no clear match for the repressor domain^31^. These findings indicate that while non-DNA-binding domains have diverged, the zinc finger-mediated DNA-binding capacity of SP1 is strongly conserved. This is consistent with the earlier study by Lin et al., which demonstrated DNA-binding and transcriptional activation capabilities of purified zebrafish Sp1^23^.

### Temporal dynamics and organ-specific localization of *Sp1* during zebrafish organogenesis

To investigate the temporal and spatial expression patterns of *sp1* during zebrafish development, we analyzed mRNA levels across key embryonic stages. Embryos were collected at the 2–4 cell, 128-cell, 1k-cell, dome (4.3 hours post-fertilization [hpf]), shield (6 hpf), bud (10 hpf), and then at daily intervals from day 1 to 5 days post-fertilization (dpf). Quantitative transcript analysis showed that *sp1* is maternally deposited and showed high abundance during the early cleavage stages, up to 1K cell stage (Figure 2A). Expression declined sharply at the dome stage and progressively decreased through gastrulation and larval development. These findings were supported by transcriptomic data from the EMBL-EBI Expression Atlas^32^, which also revealed high maternal expression followed by reduced zygotic levels, with minor timing differences likely due to methodological and normalization variations (Figure 2B). Further, we looked into the publicly available proteomics data^33^ for the presence of Sp1 protein, at least confirming the relatively higher amount of Sp1 protein during early stages of zebrafish development (Figure 2C).

**Figure 2.**
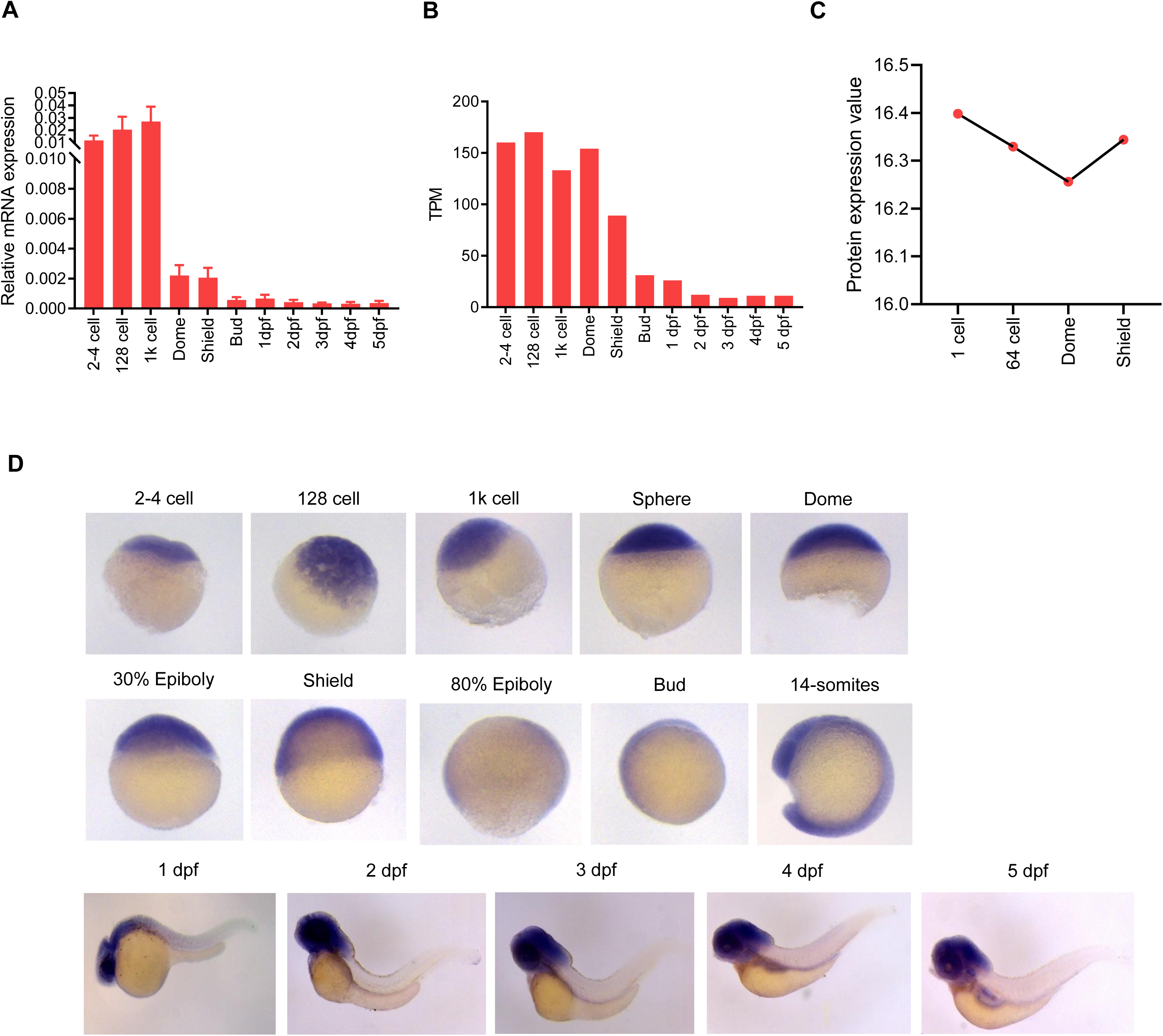
Expression dynamics of *sp1* during zebrafish development. **A.** Expression of *sp1* transcript during zebrafish development using qRT-PCR. **B.** Expression of *sp1* transcript using publicly available stagewise mRNA-seq data from the EBI-EMBL database. **C.** Expression of Sp1 protein using publicly available stagewise proteomics data. **D.** Expression and localisation of *sp1* transcript during zebrafish development using WISH. Lateral views with the anterior on top for early stages (2-4 cell to Shield) and to the left for late stages (Bud to 5 dpf) of development.

Whole mount in situ hybridization (WISH) further confirmed these patterns. Early embryos displayed robust, ubiquitous *sp1* expression, paralleling the transcript data (Figure 2D), consistent with the widespread expression reported in mouse embryos^34^. By contrast, later stages showed localized *sp1* expression in the anterior region (from 1 dpf) and the gastrointestinal tract (from 4 dpf), suggesting roles in anterior structure formation and gut development. Given that human SP1 is implicated in neuronal function and colorectal cancer progression, these observations point to the conserved developmental roles of SP1 across species^34–36^.

### Depletion of Sp1 results in defects in larval stages

To investigate the functional requirement of *sp1*, we performed knockdown experiments using a translation-blocking morpholino (MO). Wild-type embryos were injected with increasing doses (2, 4, 8, and 16 ng) to establish an effective range, with a standard control MO as a negative control (Suppl Figure S2). Although *sp1* is maternally deposited and expressed from the 2-cell stage onward, no visible abnormalities were observed at early stages. Developmental defects first became evident at 1 dpf (Suppl Figures S3A). Treatment with *sp1* MO consistently induced phenotypes across doses: 2 ng injections resulted in variable effects, while nearly all embryos injected with 4 ng showed deformities. At 8–16 ng, lethality was high, with limited variation in phenotypes. Based on this, a dosage of 2 ng was selected for further analyses (Suppl Figures S3B).

Survival assay revealed that *sp1* depletion reduced larval viability, resulting in complete lethality by 10 dpf (Figure 3A). MO-injected embryos frequently displayed cardiac arrest at 24 hpf. Gross heart malformations and anterior deformities were visible by 72 hpf (Suppl Figures S3A). To complement transient knockdown with genetic knockouts, we attempted CRISPR/Cas9-mediated deletion of *sp1*, however, no discernible phenotypes were observed in F1 embryos. This may be attributed to genetic compensation by one or more Sp family paralogs, a phenomenon commonly reported in zebrafish knockout models. Therefore, to confirm the specificity of the morpholino-induced phenotypes we co-injected a morpholino-resistant mutant form of *sp1* mRNA, which effectively rescued the observed defects (Figure 3B). Given the established involvement of human SP1 in cell cycle control and proliferation, we sought to determine whether a similar function is conserved in zebrafish. To assess the impact of *sp1* depletion on proliferation, we examined the expression of proliferation markers *mki67* and *myca*. Additionally, because *sp1* showed enrichment in the gastrointestinal system, we also assessed endodermal markers, *sox17* and *casanova*. Knockdown of *sp1* led to transcriptional alterations in both proliferation markers and endodermal regulators, supporting its functional specificity (Figure 3C&D). Interestingly, while no visible defects were detected before 1 dpf, molecular changes were already evident at 10 hpf. Conversely, by 5 dpf, when gross deformities were apparent, expression of the tested markers was no longer altered. This suggests that Sp1 exerts critical effects during early lineage specification and cell division, with continued downstream consequences that manifest morphologically at later stages.

**Figure 3.**
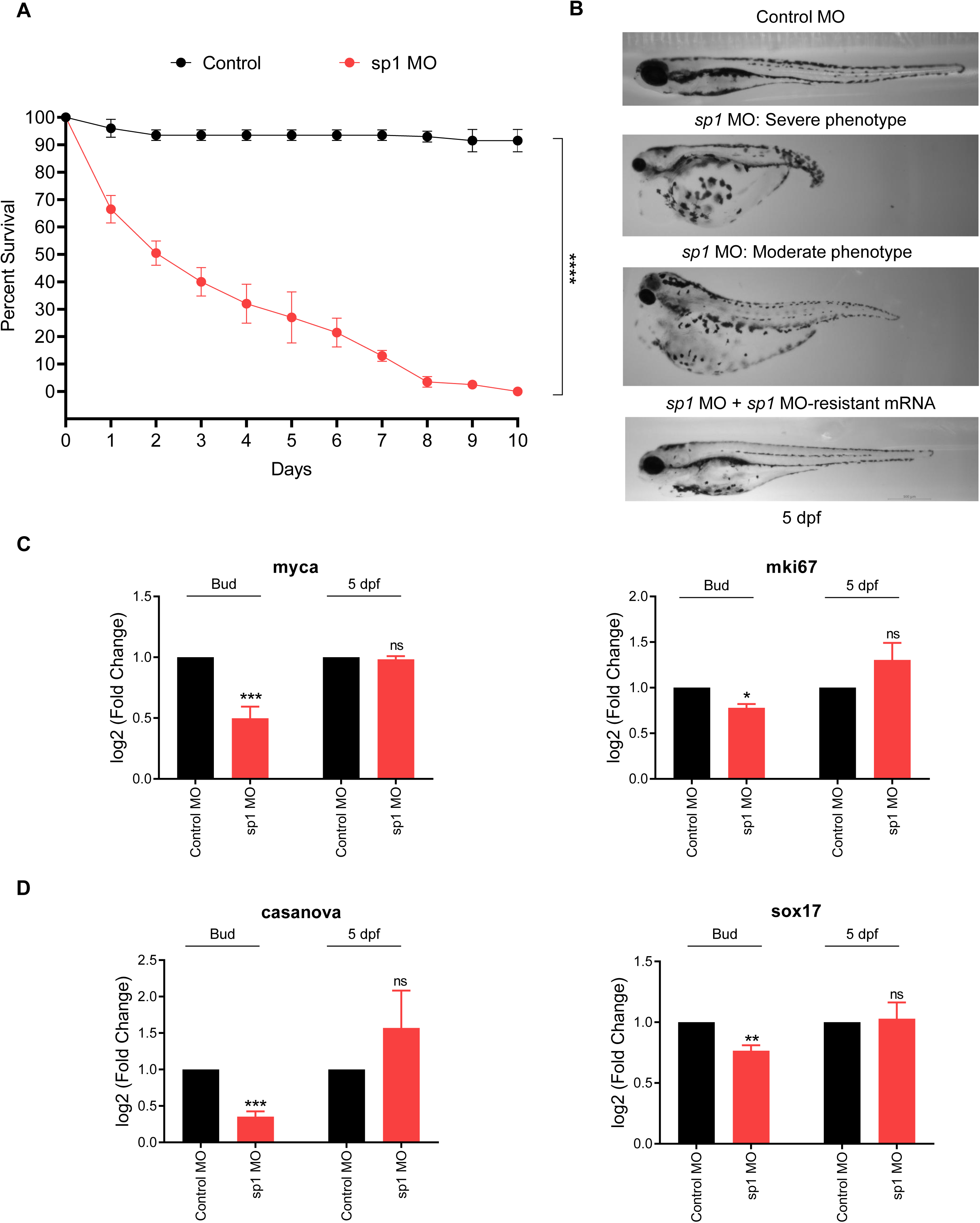
Sp1 depletion during zebrafish development. **A.** Survival analysis of zebrafish upon Sp1 depletion using the morpholino. The log-rank (Mantel–Cox) test was used to calculate significance. **B.** Morphology of 5 dpf larvae upon injection of control morpholino, *sp1* morpholino (severe and moderate deformities) and co-injection of *sp1* MO + *sp1* MO-resistant mRNA. Lateral views with the anterior to the left. Scale bar: 100µm; Magnification: 4X. **C.** Expression of proliferation markers *myca* and *mki67* using qRT-PCR in control and Sp1-depletion at 10 hpf (Bud) and 5 dpf. **D.** Expression of endodermal markers *sox17* and *casanova* using qRT-PCR in control and Sp1-depletion at 10 hpf (Bud) and 5 dpf. Unpaired t-test was used to calculate significance for panels C&D.

### Sp1 regulates fundamental cellular processes

To advance our understanding of the regulatory roles of Sp1, we conducted transcriptome analysis in Sp1-depleted zebrafish embryos using MO. Given the persistence of morpholino effects until 5 dpf and the observed defects in cranial and gastrointestinal regions, we selected three developmental stages for analysis: 10 hpf (Bud stage), 19 hpf (20 somite stage), and 3 dpf (Protruding-mouth stage) (Figure 4A). The largest number of differentially expressed genes (DEGs) was detected at 3 dpf (1701), followed by 19 hpf (1253) and 10 hpf (1111) (Figure 4B), underscoring the importance of Sp1 throughout development. Minimal overlap among dysregulated genes indicated stage-specific regulatory roles, though 10 hpf and 19 hpf shared a higher degree of overlap (Figure 4C&D).

**Figure 4.**
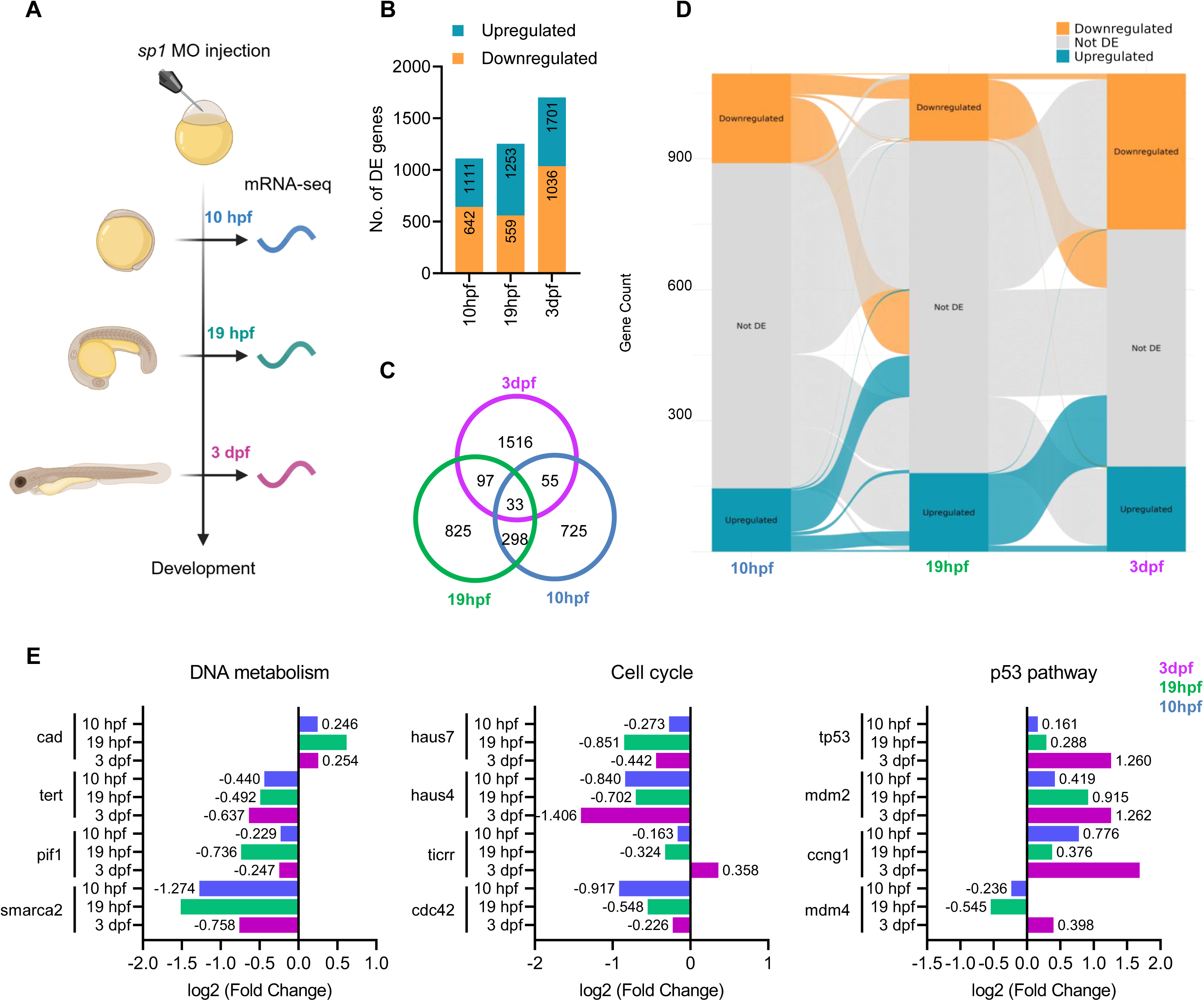
Transcriptome analysis upon Sp1 depletion. **A.** Schematic showing the stages used for transcriptome analysis upon Sp1 depletion. **B.** Number of differentially regulated (DE) genes for Bud (10 hpf), 20-somites stage (19 hpf) and protruding mouth stage (3dpf). **C.** Overlap between the differentially expressed genes in all three stages. **D.** Alluvial plot to show the differences in the sets of genes showing differential expression at the three stages chosen for transcriptome analysis upon Sp1 depletion. **E.** Expression change (log2 fold change) for marker genes for DNA metabolism (left), Cell cycle (middle) and p53 pathway (right) upon Sp1 depletion. Stages are colour-coded according to the key on the extreme right.

Gene ontology analysis of differentially expressed genes following Sp1 knockdown revealed dynamic, stage-specific enrichment patterns (Suppl Figure S4). Key processes showing downregulation include neurodevelopmental processes like visual perception, synaptic signaling, chromatin organization, DNA replication, cell-cycle regulation, pigment granule organization, vesicle transport, mitochondrial transport, and intrinsic apoptotic pathways. Enriched categories among the processes showing upregulation include regeneration, muscle cell development, immune response regulation, cilium assembly, metabolic processes, ventricular trabecula morphogenesis, ion transport, and multicellular organism biosynthesis. Upregulated genes at 3 dpf also included those involved in wound repair and p53 signaling (Suppl Figure S4A&B). Importantly, across all three stages, we identified a subset of consistently dysregulated genes significantly enriched for DNA metabolism, cell cycle progression, and p53 signaling (Figure 4E). Notably, p53 is a known SP1 interactor in humans, suggesting a conserved regulatory axis in zebrafish^37–39^. These findings establish Sp1 as a critical regulator of fundamental cellular processes throughout embryogenesis.

### Sp1 is essential for the development of multiple organ systems

Given the enrichment of *sp1* transcripts in the head and gastrointestinal regions, we next examined its role in organ-level development. Using transcriptomic datasets, we compared DEGs with curated marker genes for the skeletal (bone and cartilage), nervous (brain, spinal cord, retina), gastrointestinal (pharynx, liver, intestine, pancreas), and cardiovascular (cardiomyocytes, endothelial cells) systems^40^. A large proportion of genes associated with these systems were dysregulated upon *sp1* depletion (Figure 5A).

**Figure 5.**
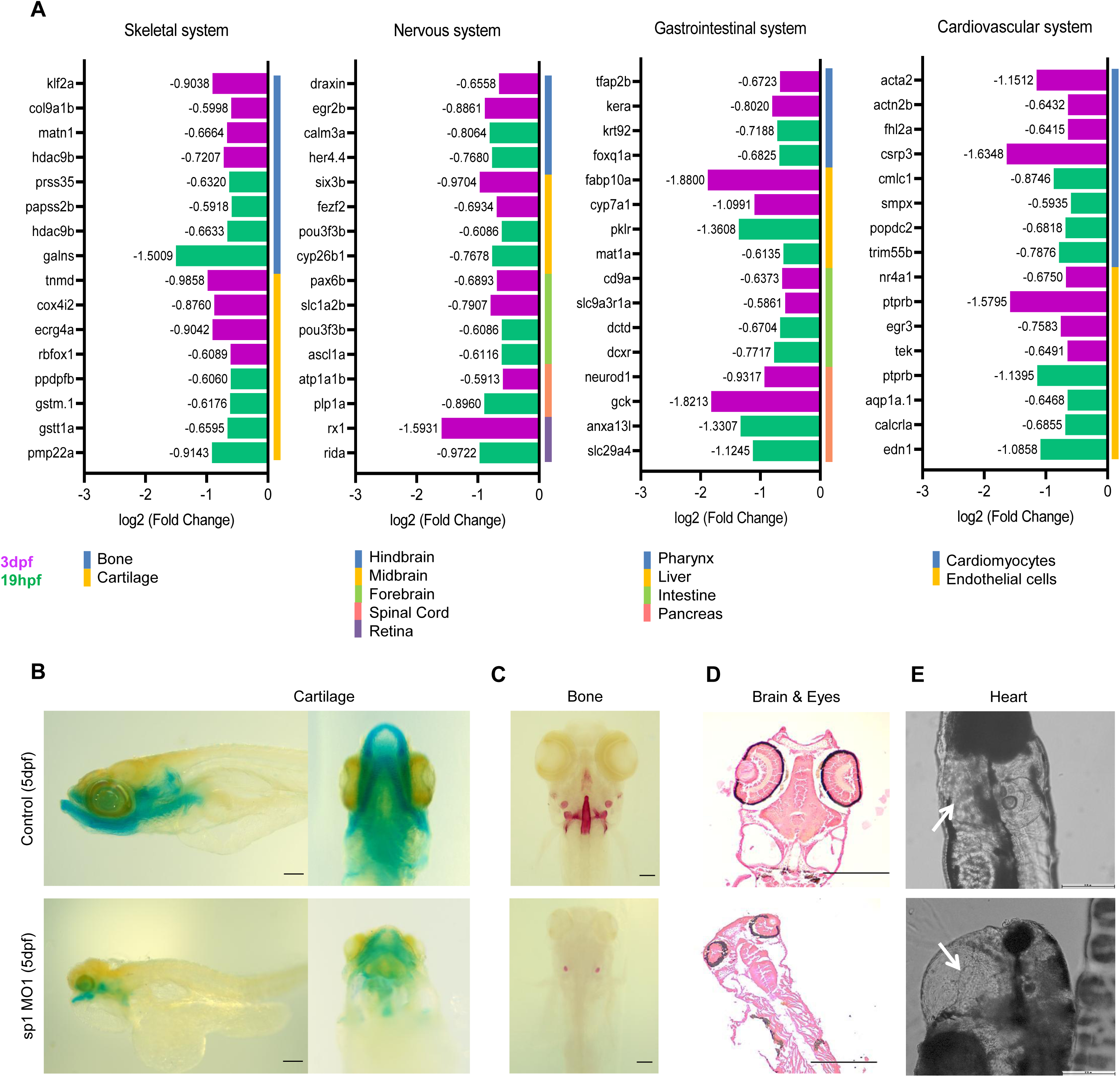
Perturbation of Sp1 causes multiple developmental defects in larvae. **A.** Expression change (log2 fold change) in the genes enriched for specific organ systems, namely (left to right) skeletal system, nervous system, gastrointestinal system and cardiovascular system. Stages are colour-coded according to the key in the lower left corner. Organs/tissues for each system are color-coded according to the key below each graph. **B.** Alcian blue staining to show cartilage development in control and Sp1-depleted larvae at 5 dpf. The left panel shows lateral view with anterior to the left. The right panel shows a ventral view with anterior on the top. Scale bar: 100µm. **C.** Alizarin staining to show bone development in control and Sp1-depleted larvae at 5 dpf. Dorsal view with anterior on the top. Scale bar: 100µm. **D.** H&E staining to show brain and eye development in control and Sp1-depleted larvae at 5 dpf. Dorsal view of an orthogonal section with anterior on the top. Scale bar: 250µm. **E.** Brightfield image to show heart development in control and Sp1-depleted larvae at 5 dpf. Lateral view with anterior on the top. Scale bar: 250µm.

To further validate these findings, we conducted a series of phenotypic assays. To assess Sp1 function in skeletal development, we performed Alcian Blue and Alizarin Red staining at 5 dpf to visualize cartilage extracellular matrix and calcified bone matrix, respectively. At this stage, multiple craniofacial cartilages are normally well formed and easily identifiable, although full bone calcification may require up to a month. Nevertheless, several bone elements are detectable at 5 dpf. As expected, control larvae displayed distinct cartilage and early-forming bone structures in the head region. In contrast, sp1 morphants exhibited severe abnormalities, including markedly underdeveloped cartilages that failed to reach appropriate size and morphology, along with the complete absence of several bone elements (Figure 5B & 5C). For the nervous system, gross morphological examination revealed a pronounced reduction in eye size and other anterior structures. Histological analysis using Hematoxylin and Eosin (H&E) staining confirmed these defects, demonstrating ocular hypoplasia along with reduced brain volume (Figure 5D). Cardiovascular phenotyping revealed profound disruption of cardiac morphology. sp1 morphants displayed a poorly differentiated linear cardiac tube, characterized by weak, irregular, and low-frequency contractions lacking clear atrioventricular coordination (Figure 5E). Collectively, these phenotypic analyses demonstrate that Sp1 is essential for the proper development of multiple organ systems in zebrafish. While human SP1 is known to broadly influence organogenesis owing to its ubiquitous expression and roles in processes such as proliferation and survival, zebrafish Sp1 shows early widespread expression that later becomes restricted to anterior and gastrointestinal regions—consistent with its broad developmental importance. Together, the transcriptomic and phenotypic results underscore Sp1 as a critical regulator of both cellular homeostasis and tissue-specific development in zebrafish.

### Sp1 is required for the formation and differentiation of the gastrointestinal system

To further investigate the role of *sp1* in gastrointestinal development, we employed the *Tg(sox17:GFP)* zebrafish line, which labels the endoderm and developing gut. By 5 dpf, when gut morphogenesis is typically complete^42^, *sp1*-depleted larvae displayed pronounced structural malformations (Figure 6A). Alcian blue staining revealed a substantial reduction in goblet cells, particularly in the posterior gut (Figure 6B), while histological analysis of transverse sections showed markedly smaller guts lacking the characteristic finger-like epithelial projections, indicative of impaired epithelial maturation (Figure 6C). This observation corroborates our earlier finding that *sp1* depletion disrupts the expression of endodermal markers, *sox17 and casanova*, during early development.

**Figure 6.**
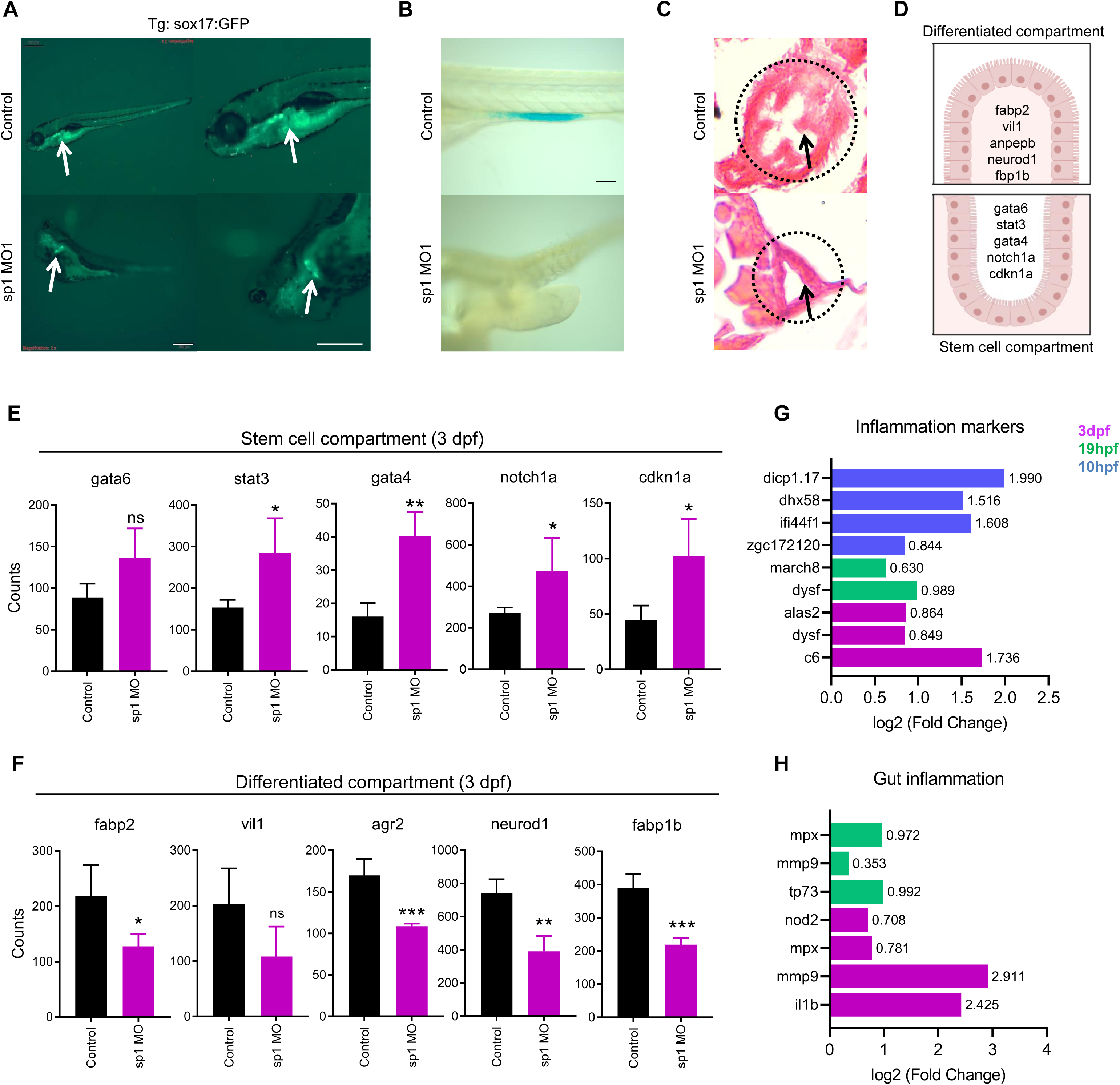
Sp1 regulates the development of the gastrointestinal system. **A.** Fluorescent image of Tg:*sox17*:GFP line to show gut development in control and Sp1-depleted larvae at 5 dpf. Lateral view with anterior to the left. The right panel is the magnified view from the left panel. Scale bar: 500µm; Magnification: 2X. **B.** Alcian blue staining to show gut-associated goblet cells in control and Sp1-depleted larvae at 5 dpf. Lateral view with anterior to the left. Scale bar: 100µm. **C.** H&E staining of a transverse section to show gut epithelium in control and Sp1-depleted larvae at 5 dpf. Scale bar: 250µm. **D.** Schematic to show genes associated with differentiated compartment (top panel) and stem-cell compartment (lower panel) of zebrafish gut. **E.** Expression of stem-cell compartment markers in control and Sp1-depleted larvae at 3 dpf. Unpaired t-test was used to calculate significance. **F.** Expression of differentiated compartment markers in control and Sp1-depleted larvae at 3 dpf. Unpaired t-test was used to calculate significance. **G.** Expression change of general inflammation markers upon control and Sp1-depletion. **H.** Expression change of gut-associated inflammation markers upon control and Sp1-depletion. Stages are colour-coded according to the key on the extreme right.

To better understand the epithelial defects and reduced goblet cell numbers, we examined markers of stem and differentiated cell populations (Figure 6D). Unlike mammals, zebrafish lack the crypt-villus structures but possess finger-like projections into the lumen where the differentiated cells reside, with stem cells localized at their base. Interestingly, while human SP1 has been implicated in maintaining stemness in colorectal cancer cells^43^, *sp1* depletion in zebrafish led to the opposite trend: stem cell-associated markers (*gata6* and *stat3*) were upregulated, whereas differentiation markers (*fabp2* and *vil1*) were significantly downregulated at 5 dpf (Suppl Figure S5). These findings were consistent with the transcriptome profile from 3 dpf, which also showed elevated stem cell marker (*gata6, stat3, gata4, notch1a* and *cdkn1a*) expression and reduced differentiation markers (*fabp2, vil1, agr2, neurod1* and *fabp1b*) (Figure 6E&F). This divergence from mammalian systems suggests a developmental stage or context-specific role for Sp1 in zebrafish, or possibly compensatory feedback triggered by morpholino-mediated depletion. Given the observed expansion of the stem cell compartment, we next investigated whether the inflammatory signaling contributed to this phenotype. Indeed, *sp1*-depleted larvae showed increased expression of both general inflammatory genes (Figure 6G) and gut-specific inflammatory markers (Figure 6H). These findings suggest that inflammation-associated cues may exacerbate epithelial defects and reinforce stem cell expansion in the absence of *sp1*.

## Discussion

This study aimed to elucidate the function of Sp1 during zebrafish development. Previous reports demonstrated that zebrafish Sp1 shares both sequence homology and DNA-binding properties with its human counterpart^23^. Our phylogenetic analysis confirmed the evolutionary conservation of SP1 across invertebrates and vertebrates, with strong conservation of the zinc finger DNA-binding domain. Sequence comparisons further revealed that zebrafish Sp1 retains nearly all domains found in human SP1, particularly the three canonical zinc finger motifs. However, the repressor domain was not identifiable, and Domain D exhibited the lowest sequence similarity among predicted domains. Interestingly, the repressor domain in human SP1 mediates transcriptional repression^44^, suggesting that ancestral Sp1 proteins may have primarily functioned as transcriptional activators, with repressive properties evolving later through domain acquisition. Similarly, the ability to oligomerize and perform synergistic activation of transcription may have been acquired during evolution. Although these hypotheses require experimental validation, they illustrate how comparative analyses across species can illuminate the evolutionary progression and diversification of transcription factor functions.

Expression profiling revealed a strong maternal contribution of Sp1 transcripts, which declined around the maternal-to-zygotic transition and later became spatially restricted to the anterior region and developing gut. This pattern aligns with the evolutionarily conserved role of Sp1 in early cellular regulation and indicates emerging organ-specific specialization during zebrafish organogenesis. Transcriptomic and phenotypic analyses further reinforced this dual functionality, sp1 depletion disrupted fundamental cellular processes, including cell cycle progression, DNA metabolism, and apoptosis, across developmental stages, while concurrently impairing organ-level programs related to brain, eye, heart, skeletal, and gastrointestinal development. This dual role mirrors the functional versatility of SP1 in mammalian systems, where it governs both ubiquitous housekeeping genes and tissue-specific developmental pathways^45,46^. Moreover, the overlap between zebrafish sp1 targets and known human SP1 interactors, such as p53, underscores the conservation of regulatory networks across vertebrate evolution^37^.

A major focus of this study was the gastrointestinal system, where Sp1 expression became progressively enriched at later developmental stages. Loss of Sp1 led to impaired epithelial maturation, reduced goblet cell numbers, and disrupted the balance between stem and differentiated cell populations. Interestingly, while human SP1 has been reported to promote stemness in colorectal cancer cells, our findings in zebrafish revealed an apparent increase in stem cell-associated markers alongside decreased differentiation markers upon *sp1* depletion^43^. This contrasting outcome may reflect stage-specific roles of Sp1, compensatory signaling mechanisms, or fundamental differences between developmental and disease contexts. Additionally, the observed elevation in inflammatory marker expression suggests that inflammation-driven feedback could contribute to the expansion of stem cell populations in sp1-deficient larvae. Collectively, these findings establish Sp1 as a crucial regulator of gut development and provide an entry point to investigate its crosstalk with major signaling pathways, including Wnt and BMP, that are central to both developmental processes and cancer biology^47^.

Comparison with mammalian systems strengthens the biological relevance of these findings. In mice, Sp1 deletion leads to lethality around embryonic day 11, with defects in multiple organ systems, consistent with the pleiotropic effects observed here in zebrafish^48^. Yet the delayed onset of overt phenotypes in zebrafish morphants until 24 hpf suggests that early embryonic events may be sustained by maternal Sp1 contribution or functional redundancy, with defects becoming apparent only during organogenesis when zygotic regulation predominates. This feature, together with the external development and accessibility of zebrafish embryos, underscores the utility of this system for dissecting stage– and tissue-specific Sp1 functions that may be masked in mammalian models.

Future directions should prioritize approaches beyond transcriptomics to direct mechanistic studies, including global mapping of chromatin binding sites, identification of transcriptional interactors, and functional dissection of domains and post-translational modifications. Such studies will clarify how Sp1 integrates ubiquitous cellular control with lineage-specific transcriptional regulation. Equally important will be systematic analysis of potential paralog compensation, which likely contributes to the lack of phenotype in sp1 morphants at early stages, as well as the generation of zebrafish-specific antibodies for robust protein-level assays. Together, these strategies will enable deeper exploration of how Sp1 continues to reveal unanticipated and fundamental roles in vertebrate biology, despite being one of the earliest transcription factors discovered.

## Materials and Methods

### Zebrafish husbandry

Wild-type strain TU of zebrafish and transgenic *sox17*:GFP lines were maintained according to the standard zebrafish husbandry protocols under the National Facility for Gene Function in Health and Disease, IISER, Pune and Center for Integrative and Translational Research (CITRES), SNIOE, Delhi-NCR. Embryos and larvae were grown at 28 °C in 1X E3 medium (5 mM NaCl, 0.33 mM CaCl_2_, 0.33 mM MgSO_4_, 0.17 mM KCl, and 0.1% Methylene Blue) till harvesting. Bright-field images for rescue experiments were acquired using the ZEISS Axiocam 506 mono camera. All remaining bright-field and fluorescence imaging was performed using an Olympus stereo microscope.

### Phylogenetic analysis of SP1 proteins

SP1 Protein sequences were retrieved from the NCBI Protein database (https://www.ncbi.nlm.nih.gov/protein/)^49^. Orthologous sequences from species included in the NCBI ortholog set were selected for analysis. Multiple sequence alignments were performed using the MAFFT L-INS-i algorithm, and the resulting alignments were trimmed using trimAl in automatic mode^50,51^. Both steps were implemented via the NGPhylogeny.fr online server (https://ngphylogeny.fr)^52^. Phylogenetic tree construction was carried out using the Randomized Accelerated Maximum Likelihood (RAxML-NG) algorithm on the CIPRES Science Gateway server^53,54^. The best-fit model of protein sequence evolution was determined using ModelTest-NG, which identified JTT+I+G4 as the optimal model. Accordingly, tree inference was performed using the JTT+I+G4m model in RAxML-NG, which employs the JTT substitution matrix, a discrete gamma distribution with four categories using mean rate values (+Gm), maximum likelihood estimation of the proportion of invariant sites (+I), and empirical amino acid frequencies (+F). To root the tree, *Branchiostoma belcheri* was designated as the outgroup. A total of 1000 bootstrap replicates were performed to assess branch support, with the analysis conducted in –-search mode. The resulting phylogenetic tree was visualized using the Interactive Tree of Life (iTOL) tool^55^. Representative animal silhouettes were obtained from PhyloPic (http://phylopic.org) and annotated onto the tree^56^.

### Protein structure prediction and comparative analysis

The predicted structure of the human SP1 protein was obtained from the AlphaFold Protein Structure Database^57,58^ (AF-P08047F1-Predicted), while the zebrafish SP1 protein structure was predicted using AlphaFold3^59^. From the five structures generated by the AlphaFold server, the most suitable structure was selected based on residue distribution in the Ramachandran plot, which indicated favorable conformations. The full-length protein structures for human and zebrafish SP1 were aligned pairwise using TM-align^60^ (https://www.rcsb.org/alignment) for structural comparison. Domain information for the human SP1 protein was extracted using residue positions obtained from the UniProt database^61^ (www.uniprot.org). For the zebrafish SP1 protein, the corresponding domain was identified by aligning the amino acid sequences of human and zebrafish SP1 proteins using BlastP^49^ (https://blast.ncbi.nlm.nih.gov/Blast.cgi), and residues from the aligned regions were used to extract the zebrafish domain structure using PyMOL (The PyMOL Molecular Graphics System, Version 1.2r3pre, Schrödinger, LLC.). The extracted domains from both species were subsequently aligned pairwise using the flexible alignment algorithm JFATCAT^62^ (https://www.rcsb.org/alignment) for structural comparison^63^.

### Morpholinos and mRNA microinjections

Morpholino for *sp1* and control morpholino were obtained from Gene Tools, LLC. Morpholinos were resuspended in nuclease-free water according to the manufacturer’s instructions. Embryos were injected with morpholinos at a 0.5 nM concentration at the 1-cell stage and grown till the indicated stage. Mutant sp1 mRNA was generated using mMESSAGE mMACHINE T7 Transcription Kit (Thermo Fisher Scientific) according to the manufacturer’s instructions. 6.25pg of mRNA was co-injected along with morpholino for performing the rescue experiments. A list of the morpholino sequences and primers used for designing the mutant mRNA is given in Supplementary Table 1.

### Survival analysis

Survival analysis was performed by injecting the standardized dose of morpholinos into 50 embryos per replicate (three replicates in total). Embryos were maintained under standard conditions (28 °C in 1X E3 medium), monitored and cleaned daily, and mortality was recorded up to 10 dpf. The absence of a heartbeat was used as the criterion for mortality. Statistical significance was assessed using the log-rank (Mantel–Cox) test.

### RNA isolation, RT-qPCR and analysis

Twenty dechorinated larvae (or embryos) per replicate were lysed in 1 mL of RNAiso Plus (Takara), followed by a standard extraction protocol. Briefly, the cells were homogenised using a Pellet Pestle Motor (Kontes). 200 μL chloroform was added, followed by centrifugation at 12000 rpm at 4°C for 15 mins. The aqueous layer was collected in a fresh tube, followed by another chloroform wash. Precipitation was performed by adding 10% of 3M sodium acetate solution and an equal volume of 100% isopropanol containing Glycoblue coprecipitant (Invitrogen) and incubating at –20°C overnight. RNA was pelleted at 12000 x g at 4°C for 1 hour, followed by two washes of 75% ethanol at 12000 x g, room temperature (RT), for 5 minutes. cDNA synthesis was performed using the PrimeScript RT Reagent Kit (Takara) according to the manufacturer’s protocol. qRT-PCR was performed using TB Green Premix Ex Taq II (Takara) using the protocol provided by the manufacturer in the ABi ViiA 7 Real-Time PCR System. qRT-PCR analysis was performed by subtracting Ct values for the control gene, 18S rRNA, from the gene of interest to obtain ΔCt. ΔΔCt values were obtained by subtracting the ΔCt value for the control sample from the treated sample. Internal normalisation was performed by dividing all the ΔΔCt values by the ΔΔCt of the control sample. A list of the primers used for qPCR is provided in Supplementary Table 1.

### mRNA sequencing and analysis

RNA isolation was performed as described previously and processed for library preparation. Total RNA was resuspended in nuclease-free water (Ambion), followed by quantification using Qubit RNA HS system (Thermo Fisher Scientific) and RNA integrity determination using RNA 6000 Nano Kit on Bioanalyser 2100 (Agilent). RNA samples with RIN values greater than 8 were used for library preparation. 500 ng of total RNA was subjected to mRNA purification using NEBNext Poly(A) mRNA Magnetic Isolation Module (New England Biolabs) according to the manufacturer’s instructions. The purified mRNA was used for library preparation using the NEBNext Ultra II RNA Library Prep Kit for Illumina (New England Biolabs) using the protocol provided in the kit. The final libraries were purified using the HighPrep PCR Clean-up System (MagBio Genomics, USA). The libraries were quantified using the Qubit 1X HS DNA system (Thermo Fisher Scientific). All the libraries were pooled in equimolar ratios and subjected to 75 bp PE chemistry on the Nextseq550 sequencer (Illumina), aiming at least 30 million reads per sample. Sequencing data was processed as described in Pradhan et al 2021^64^. Briefly, the bcl files obtained from sequencing were demultiplexed and converted to fastq files for further analysis. The sequencing reads were trimmed using Trimmomatic-0.39 and aligned to the DanRer10 genome assembly for zebrafish using the HISAT2 alignment program^65,66^. Gene feature counts were calculated using the FeatureCounts package from Rsubread^67^. EdgeR was used to perform differential expression analysis for 3-4 replicates per condition^68^. Metascape was used for the Gene Ontology analysis^69^.

### Riboprobe synthesis for whole-mount *in situ* hybridisation

Primers for the riboprobes were designed against zebrafish sp1 using NCBI Primer Blast^49^. The SP6 and T7 promoter sequences were added to the forward and reverse primers, respectively. cDNA was prepared using RNA extracted from the 128-cell stage using the High-Capacity Reverse Transcription kit (Applied Biosystems). The template for riboprobe synthesis was prepared by PCR using GXL Polymerase (Takara). The template was purified using the Monarch Gel extraction kit (New England Biolabs). The sense and antisense riboprobes were then prepared at 37°C using the SP6 and T7 DIG RNA Labelling kit (Roche), respectively. The riboprobes were purified using the Purelink PCR purification kit (Invitrogen). The purified riboprobes were quantified using Nanodrop and diluted with HM+(50% v/v Deionised formamide,5X SSC, 50 mg/mL Heparin stock, 0.1% Tween 20, 0.5 mg/mL RNase-free tRNA, adjusted to pH 6 using 1M Citric acid) to 30ng/ul and stored at –80 °C. A list of the primers used for probe synthesis is provided in Supplementary Table 1.

### Whole-mount in situ hybridisation (WISH)

WISH was performed as described in Thisse and Thisse, 2008^70^. Wild-type embryos were fixed using 4% paraformaldehyde (PFA) at various stages and incubated at 4°C overnight with gentle rocking. The embryos were then dechorionated with watchmakers’ forceps and dehydrated in 100% methanol. The embryos were stored at –20°C until the start of WISH. On the first day, the embryos were then downgraded using various dilutions of methanol in 1X Phosphate Buffered Saline with 0.1%v/v Tween 20 (PBST: 1X PBS: 137 mM NaCl, 2.7 mM KCl, 10mM Na_2_HPO_4_, 1.8 mM KH_2_PO_4_, adjusted to pH 7.4) (75%, 50%, 25%). After a final wash with PBST, the embryos were permeabilised using 5 μg/μl of Proteinase K in PBST for the following periods in accordance with the stage: Till bud-No treatment; 14-somite-2 minutes; 24 hpf-10 minutes; 48 hpf and above-30 minutes. After Proteinase K treatment, the embryos were refixed in 4% PFA for 20 minutes at room temperature. After PBST washes, the embryos were prehybridised in HM+ for 4 hours at 70°C. The pre-hybridised embryos were then incubated with 400-500 ng of riboprobe for 12-16 hours at 70°C. The embryos were then washed according to the protocol. The embryos were then incubated in blocking buffer (2% v/v goat serum, 2mg/mL BSA, in PBST) for 4 hours at room temperature. Following this, the embryos were incubated in the anti-DIG-AP Fab fragments (1:5000, Roche) at 4°C for 12-16 hours. Embryos were then washed with PBST and then alkaline Tris buffer. Embryos were then stained with BM Purple (Roche) until appropriate staining was observed. The staining was stopped using the stop solution (1X PBS pH 5.5, 1 mM EDTA pH 8.0, 0.1% v/v Tween 20). Embryos were stored at 4°C until imaging. Embryos were mounted in agarose moulds or on slides using 2% methylcellulose (for larval stages) and imaged using the Leica DFC450C microscope.

### Alcian blue and Alizarin red staining

Alcian blue and alizarin red were used to visualize developing cartilage/goblet cells of the intestinal epithelium and bones, respectively. Briefly, uninjected controls and *sp1* MO-injected 5 dpf larvae were euthanised using the cold shock method and fixed overnight in 4% PFA at 4°C on an end-to-end rotor. Larvae were then washed with successive 1X PBS washes, followed by dehydration with increasing concentration of ethanol and stored in 100% ethanol. For alcian blue staining, larvae were rehydrated to 70% ethanol and washed with 70% ethanol containing 0.1% HCl for 30 mins. The larvae were then stained using alcian blue (Sigma Aldrich) overnight at RT using 0.1 % alcian blue prepared in ethanol: acetic acid mixture (pH ∼2) with 0.1% Triton X-100. The larvae were then washed using 70% ethanol, followed by depigmentation using a 1:1 ratio of 2% KOH and 3% H_2_O_2_ for 20 mins at RT. The larvae were then rehydrated, followed by clearing using increasing concentrations of glycerol in 0.25% KOH. Larvae were stored in a 70% glycerol solution in 0.1% KOH storage solution until imaging. For alizarin staining, larvae were dehydrated to 70% ethanol and depigmented using the protocol mentioned above. This was followed by a 0.5 % KOH wash and staining with 0.01% Alizarin red (Sigma Aldrich) prepared in autoclaved MilliQ water (pH ∼ 4) for 2 h at RT. The larvae were then washed with 0.5 % KOH and immediately proceeded for imaging. At the time of imaging, both alcian blue and alizarin red-stained larvae were mounted with the help of low-melting agarose on a 2% agarose pad and imaged using a stereomicroscope (Leica S8 APO).

### Sectioning for histology

Histological preparation of uninjected control and SP1 morpholino-injected zebrafish larvae at 5 dpf was performed following the protocol described by Cooper et al., 2018^71^. Larvae were euthanized and fixed overnight at RT in 4% PFA. After fixation, specimens were washed twice with 1× PBS for 15 min each. Dehydration was carried out at RT using a graded ethanol series: 70% ethanol overnight, followed by 80% and 90% ethanol for 2 hours each, and two changes of 100% ethanol for 3 h. Dehydrated larvae were stored at 4 °C until further processing. Tissue clearing was done by incubating larvae in xylene until they appeared translucent (typically 1–2 minutes). Cleared larvae were infiltrated with molten Paraplast (Sigma-Aldrich) at 65 °C in a heated chamber: initially for 2 hours, followed by two 3-hour changes, and then overnight incubation. Larvae were oriented appropriately and embedded in paraffin blocks, which were then transferred to –20 °C for 2–3 hours to solidify. Blocks were sectioned at 10 μm thickness using a Leica HistoCore MULTICUT microtome. Sections were mounted onto 0.01% poly-L-lysine-coated slides using a 42°C water bath and dried overnight at the same temperature.

### Hematoxylin and Eosin (H&E) Staining

Slides were heated at 65 °C for 5 minutes to melt the wax and rinsed under running water. Sections were then cleared in xylene for 5 minutes and rehydrated through a descending ethanol gradient (100% to 70%, 3 mins each), followed by a wash in running tap water. For nuclear staining, slides were immersed in Delafield’s hematoxylin (Merck) for 3 minutes, rinsed in tap water, differentiated with 1% acid alcohol, and washed again with water. Counterstaining was performed with 0.5% alcoholic eosin Y for 30 seconds, followed by rinsing under running tap water. Slides were air-dried, mounted using DPX Mountant (Fisher Scientific), and imaged using a stereomicroscope (Leica S8 APO).

### Data availability

mRNA-seq raw data files have been deposited to the EMBL-EBI ArrayExpress database and are publicly available as of the date of publication under the accession number E-MTAB-16004.

## Acknowledgements

The authors are grateful to Prof. CP Heisenberg (IST Austria) for providing the Tg:*sox17*:GFP line, and Dr. Chinmoy Patra (ARI Pune, India) for the Leica DFC450C microscope used for WISH imaging. The authors thank the core facilities of IISER Pune and SNIoE Delhi-NCR, especially the IISER Pune Experimental Animal Facility (National Facility for Gene Function in Health and Disease) and the SNIoE Animal Facility (at the Center for Integrative and Translational Research) for maintaining the zebrafish strains. Figures 4A and 6D were created using BioRender.

## Author contributions

SG and AS designed the research work and wrote the manuscript with inputs from SJP and GJD. AS performed microinjections, survival assay and quantification of the phenotypes upon Sp1 depletion and prepared libraries for sequencing. SM performed preliminary phylogenetic analysis, WISH, qRT-PCR, and standardisation of MO and mRNA dosing. GJD performed the phylogenetic analysis, conducted Alcian Blue and Alizarin Red stainings, and assisted with the design and execution of the morpholino rescue experiment. SJP was involved in data interpretation and performed transcriptome analysis. PDG performed sectioning and H&E staining. All the authors have read and approved the manuscript.

## Supplementary Figure Legends

**Supplementary Figure S1.**
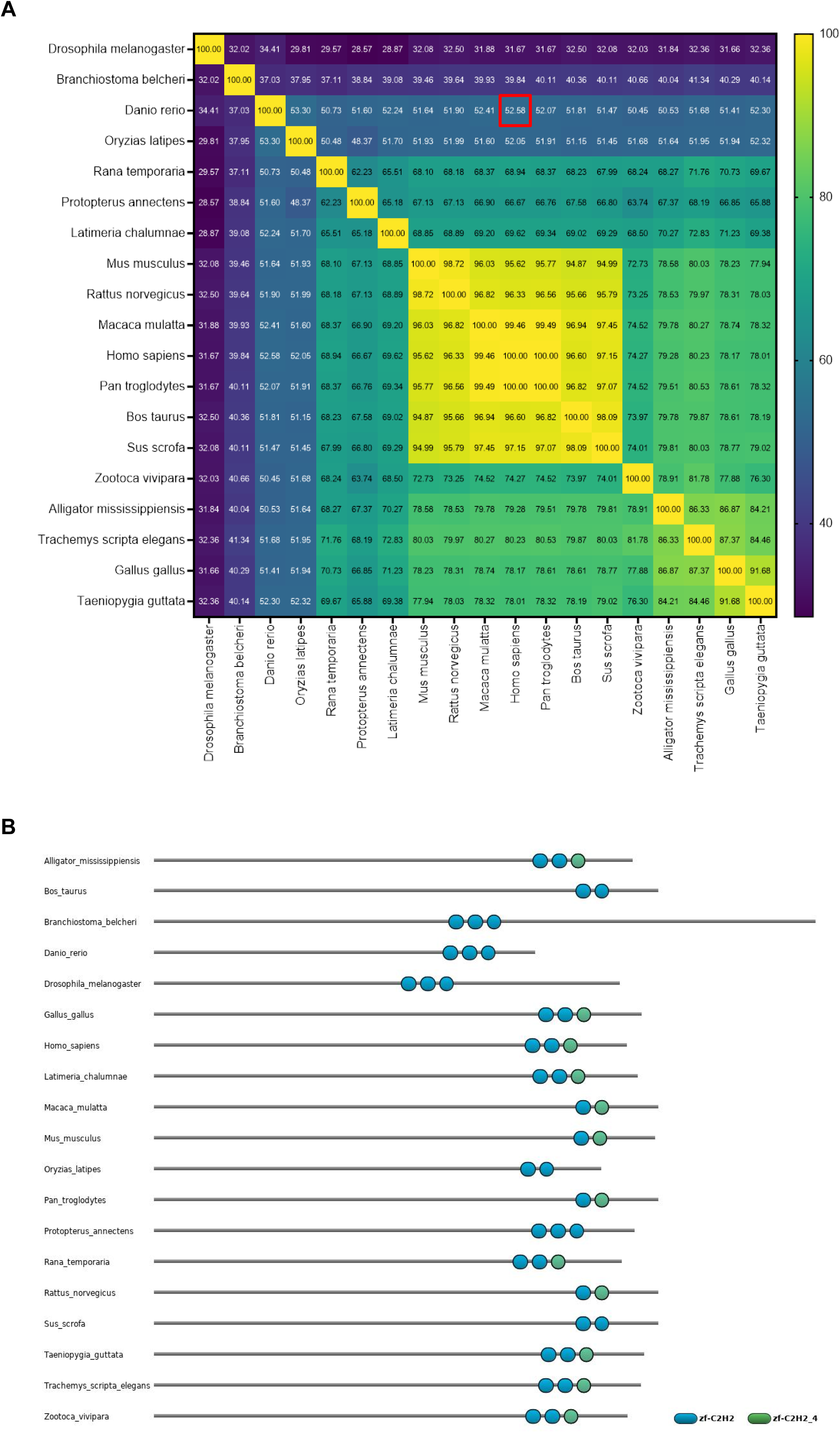
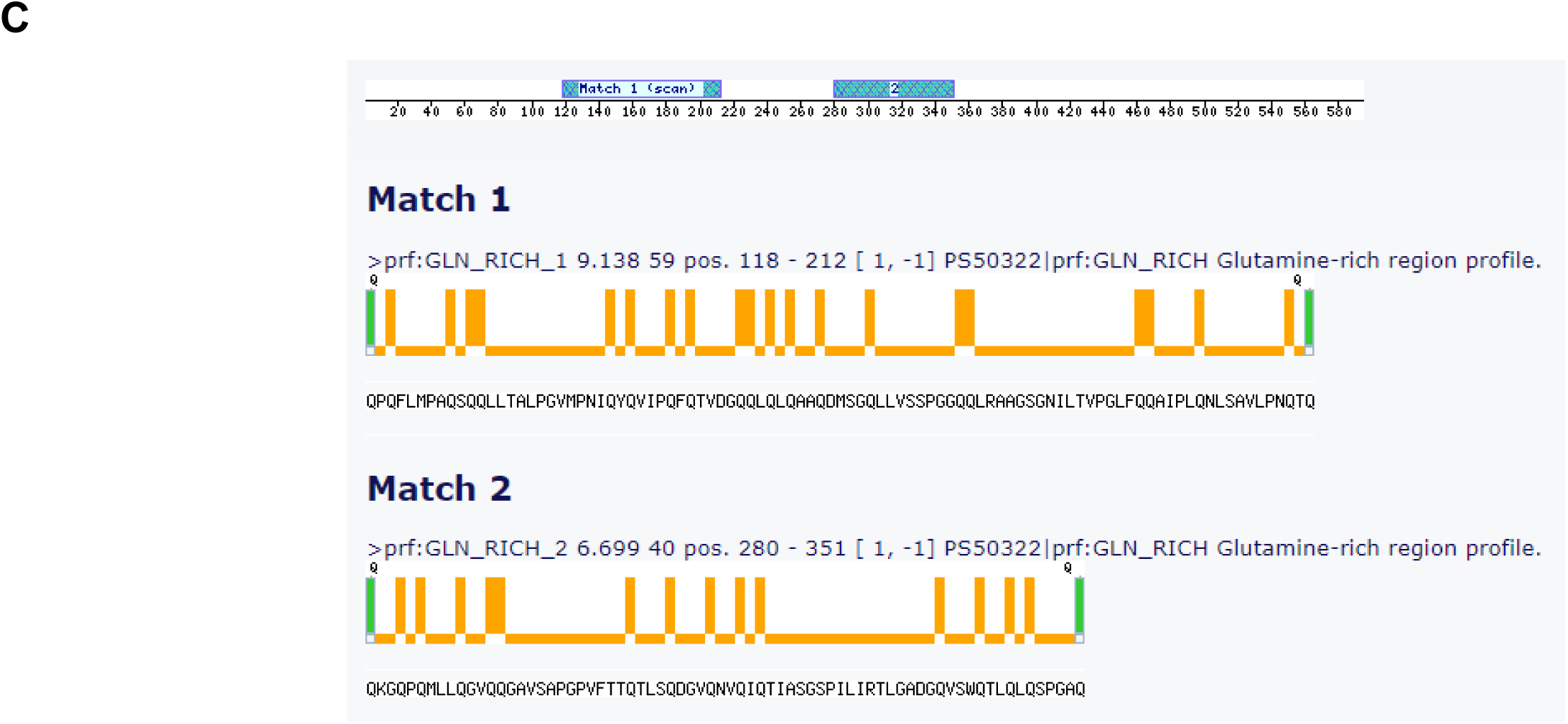
Phylogenetic analysis of SP1 orthologs: **A.** Percent identity matrix based on multiple sequence alignment of SP1 sequences generated using Clustal Omega. Red square is highlighting the similarity between human and zebrafish orthologs. **B.** Domain prediction and alignment of SP1 orthologs via HMMER algorithm using the Pfam database. **C.** Prediction for Glutamine-rich regions in zebrafish Sp1 using Motifscan.

**Supplementary Figure S2.**
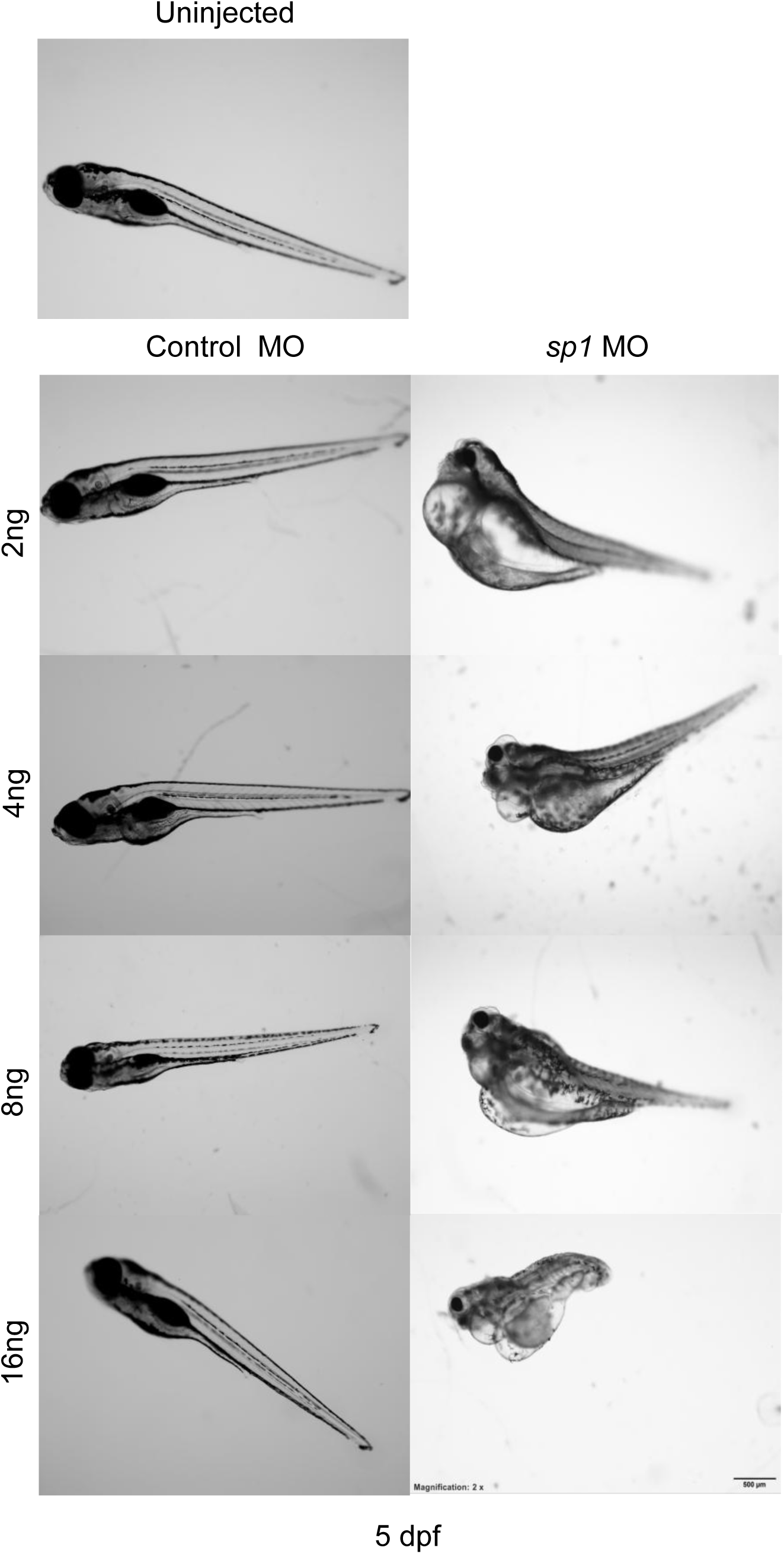
Dosage standardisation of the morpholino used in the study: Both the morpholinos were injected at a range of 2ng to 16ng and observed till 5 dpf for morphology and lethality. Lateral views with the anterior to the left. Scale bar: 500µm; Magnification: 2X.

**Supplementary S3.**
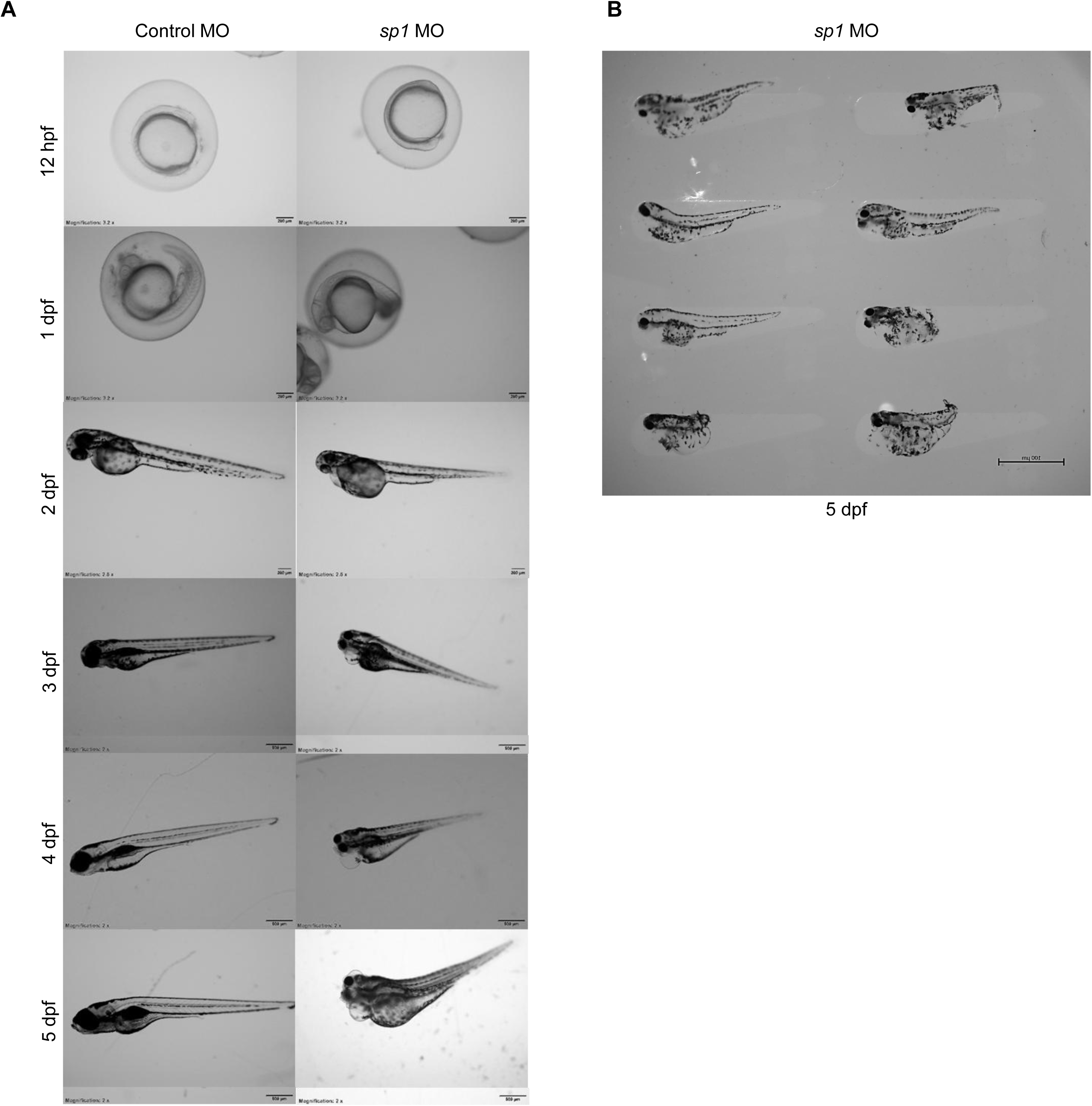
Morphology of zebrafish injected with morpholinos: **A.** Both morpholinos were injected at a dosage of 2ng. sp1 MO-injected larvae show phenotypic deformities at 5 dpf, while the control MO-injected larvae are phenotypically normal till 5 dpf. Lateral views with the anterior to the left. Scale bar: 200µm for 12 hpf and 1 dpf & 500µm for 2-5 dpf; Magnification: 3.2X for 12 hpf and 1 dpf & 2X for 2-5 dpf. **B.** Variation in the phenotypes at 5dpf observed upon sp1 depletion using MO at 2ng. Lateral views with the anterior to the left. Scale bar: 100µm; Magnification: 4X.

**Supplementary Figure S4.**
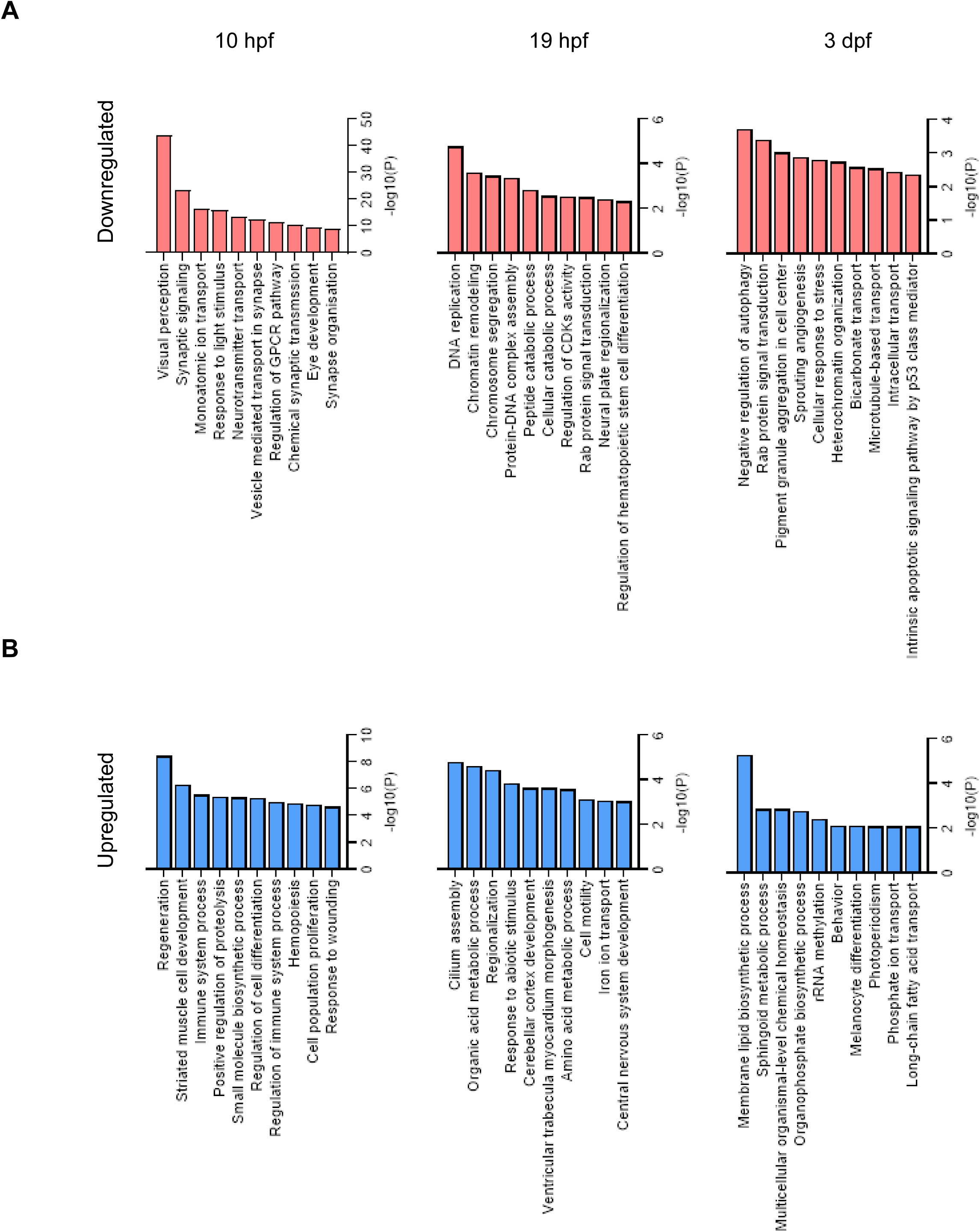
Transcriptome analysis upon Sp1 depletion. **A.** Gene ontology analysis for downregulated genes at 10 hpf, 19 hpf and 3 dpf upon Sp1 depletion. **B.** Gene ontology analysis for upregulated genes at 10 hpf, 19 hpf and 3 dpf upon Sp1 depletion. Analysis was performed using Metascape.

**Supplementary Figure S5.**
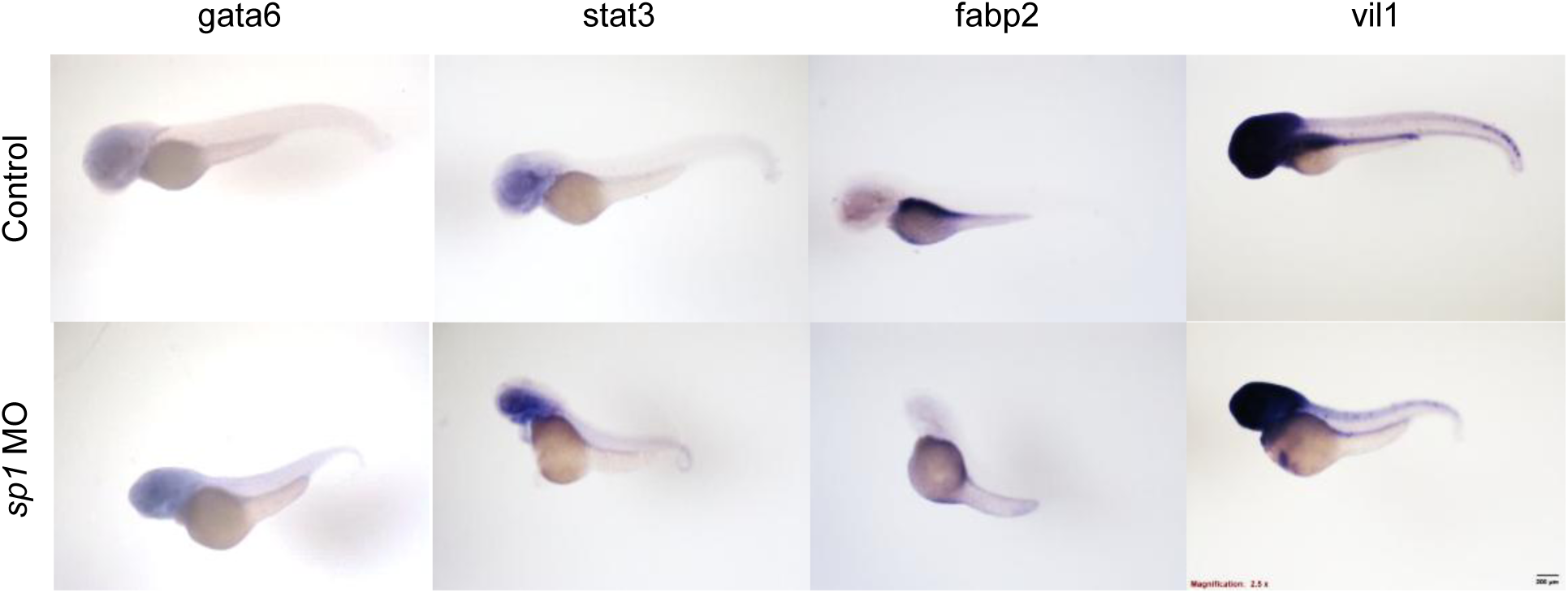
Transcriptome analysis upon Sp1 depletion. WISH to show the expression of gut-specific markers in control and Sp1-depleted at 5 dpf. Lateral views with the anterior to the left. Scale bar: 500µm; Magnification: 2X.

**Supplementary Table 1.**
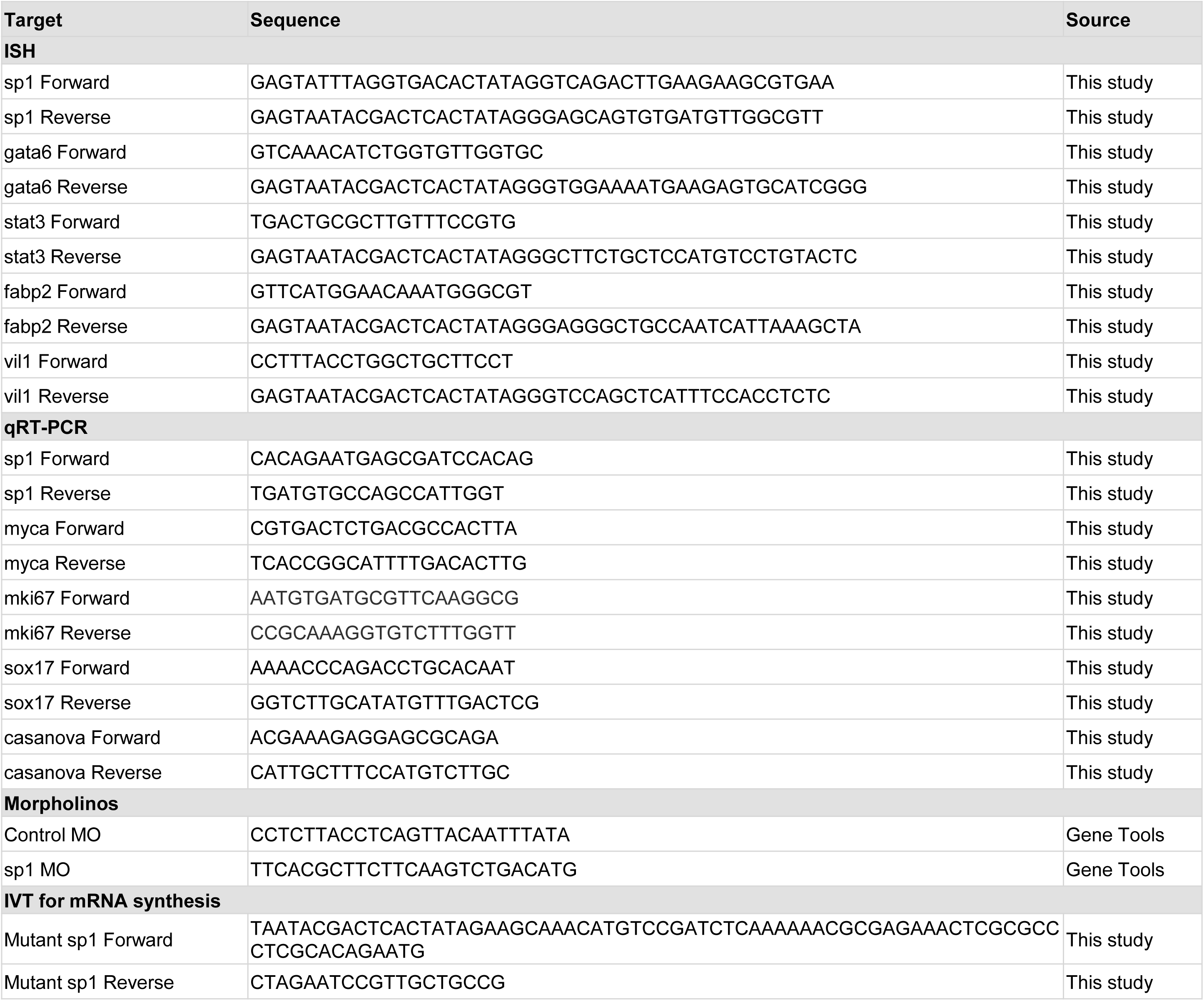
List and sequence of all the oligonucleotides used in this study: Primers used for ISH probe synthesis, qRT-PCR, morpholinos and IVT for mRNA synthesis.

